# Genome sequence of the ornamental plant *Digitalis purpurea* reveals the molecular basis of flower color and morphology variation

**DOI:** 10.1101/2024.02.14.580303

**Authors:** Jakob Maximilian Horz, Katharina Wolff, Ronja Friedhoff, Boas Pucker

## Abstract

*Digitalis purpurea* (foxglove) is a widely distributed ornamental plant. Here, we present a long read sequencing-based genome sequence of a magenta flowering *D. purpurea* plant and a corresponding prediction of gene models. The high assembly continuity is indicated by the N50 of 4.3 Mbp and the completeness is supported by discovery of about 96% complete BUSCO genes. This genomic resource paves the way for an in-depth investigation of the flower pigmentation of *D. purpurea*. Structural genes of the anthocyanin biosynthesis and the corresponding transcriptional regulators were identified. The comparison of magenta and white flowering plants revealed a large insertion in the anthocyanidin synthase gene in white flowering plants that most likely renders this gene non-functional and could explain the loss of anthocyanin pigmentation. Furthermore, we found a large insertion in the *DpTFL1/CEN* gene to be likely responsible for the development of large terminal flowers.

## Introduction

*Digitalis purpurea*, commonly known as foxglove, is a biennial plant in the Plantaginaceae family [1]. The species is native to Europe, but appears in many other parts of the world. *D. purpurea* has been used for centuries for its medicinal properties, particularly for treating heart conditions such as atrial fibrillation and congestive heart failure [2, 3]. The plant contains a variety of cardiac glycosides, with the most important being digoxin [2]. These compounds work by increasing the force and efficiency of heart contractions, which are used to treat heart failure and other cardiac conditions. *D. purpurea* and the close relative *D. lanata* are cultivated for the extraction of cardenolides [1]. Scientists have been studying the plant for many years to better understand the chemistry of the cardiac glycosides it produces and their effects on the human body. Besides its medicinal value, *D. purpurea* is also famous for beautiful flowers (**Fig. 1a**). It is a tall plant that can reach up to six feet in height and produces striking magenta flowers on tall spikes. Various *Digitalis* species display a plethora of diverse flower colors and shapes. While the plant is visually appealing, it is considered toxic due to the formation of specialized metabolites like the cardenolides. Nonetheless, *D. purpurea* is widespread as an ornamental plant. Contributing to the flower colors are anthocyanins, the product of one specific branch of the flavonoid biosynthesis (**Fig. 1b**). The substrate of the flavonoid biosynthesis is the aromatic amino acid phenylalanine, which is channeled through the general phenylpropanoid pathway and the flavonoid biosynthesis towards anthocyanins [4, 5]. Major branches of the flavonoid biosynthesis lead to colorful anthocyanins, colorless flavonols, colorless flavones, and proanthocyanidins which appear dark after oxidation. The anthocyanin biosynthesis shares several enzymatic steps with other pathways like the proanthocyanidin biosynthesis. Dihydroflavonol 4-reductase (DFR), anthocyanidin synthase (ANS), UDP-glucose dependent 3-O-anthocyanidin-glucosyltransferase (UFGT), and other decorating enzymes are often considered as the anthocyanin biosynthesis branch [4, 6] (**Fig. 1c**). A recent study revealed anthocyanin-related glutathione S-transferase (arGST) as an additional enzyme in the anthocyanin biosynthesis [7]. After biosynthesis of anthocyanins at the endoplasmic reticulum, a transport through the cytoplasm and import into the central vacuole is required for long term storage. This process involves several proteins including two tonoplast-localized transporters and a tonoplast-localized ATPase for generation of a proton gradient [5, 8, 9]. A previously postulated ‘ligandin’, that might protect anthocyanins during the transport through the cytoplasm [10], was recently characterized as an enzyme in the cyanidin biosynthesis, which makes an involvement in the anthocyanin transport less likely [7]. Besides direct transport through the cytoplasm, there are several studies that report an involvement of vesicles in the anthocyanin transport in different plant species [5, 11–14].

**Fig. 1:**
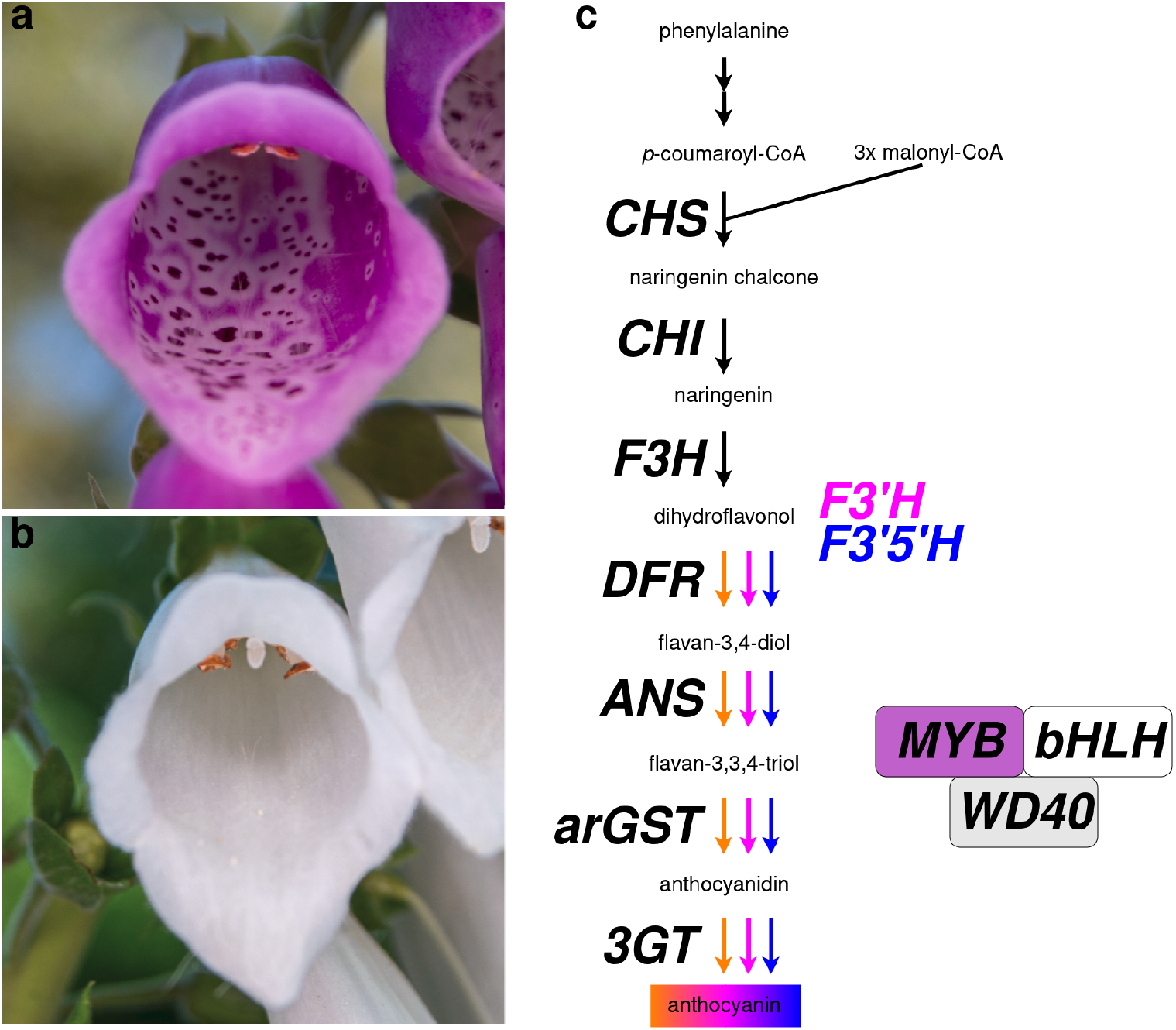
Pictures showing magenta (a) and white (b) flowers of *Digitalis purpurea* (foxglove). The anthocyanin biosynthesis is displayed as the central branch of the flavonoid biosynthesis (c). Abbreviations: CHS, chalcone synthase; CHI, chalcone isomerase; F3H, flavanone 3-hydroxylase; DFR, dihydroflavonol 4-reductase; ANS, anthocyanidin synthase; arGST, anthocyanin-related glutathione S-transferase; 3GT, UDP-glucose anthocyanidin 3-O-glucosyltransferase; MYB, Myeloblastosis protein; bHLH, basic helix-loop-helix; WD40, proposed scaffolding protein.

The pigmentation of a plant structure is not only determined by the activity of the anthocyanin biosynthesis, but also by the activity of pathways that compete for the same substrate. High activity of the flavonol biosynthesis can deplete the available substrate of the anthocyanin biosynthesis branch [15]. Subtle differences in substrate specificity of the competing enzymes,e.g. flavonol synthase (FLS) and DFR, can determine the pattern of synthesized flavonoids [16, 17]. Differences in expression of the first committed genes *DFR* and *FLS*, respectively, appear as a decisive factor in controlling the activity of the anthocyanin and flavonol biosynthesis branch [17].

Activity of different branches of the flavonoid biosynthesis is mostly controlled at the transcriptional level through regulation of gene expression by transcription factors [18–21]. Decades of research turned the regulation of the flavonoid biosynthesis into a model system for research on cis- and trans-regulatory elements in eukaryotes. The anthocyanin biosynthesis genes are activated by a complex comprising three different proteins: a R2R3-MYB protein, a bHLH protein, and a WD40 protein [21–23]. Different subgroup 6 MYBs and subgroup IIIf bHLHs can be included in this complex leading to a diverse set of different complexes which differ in their ability to activate certain target genes [24, 25]. The WD40 component is often TTG1, but LWD1 and LWD2 can also participate in complex formation [26]. An additional WRKY component (TTG2) has been suggested to specifically increase the expression of genes associated with the anthocyanin transport into the vacuole [21, 27]. Anthocyanin biosynthesis-regulating MYBs have been reported in different MYB clades. PAP1 (MYB75)/PAP2 (MYB90)/PAP3 (MYB113)/PAP4 (MYB114) are well known anthocyanin regulators in *Arabidopsis thaliana* [23, 28, 29]. While most anthocyanin regulating MYBs are activators, the incorporation of subgroup 4 MYBs into the MBW complex can result in an inhibitory function [30].

Pigmentation differences between plants of the same species are often due to genetic variations affecting the functionality or gene expression of the MYB and bHLH transcription factors [31–34]. This results in extremely low or no transcriptional activation of specific flavonoid biosynthesis genes under certain conditions. However, activation of the structural genes in the context of a different pathway is still possible [34]. Loss of function or gene loss affecting biosynthesis genes or transporters involved in the biosynthesis pathway are rare, but can explain deeper loss of the anthocyanin pigmentation in the Caryophyllales and might be irreversible [35]. Changes in flower color can affect the attraction of pollinating animals and are thus subject to selection. For example, the transition of purple blue to orange-red in *Ipomoea quamoclit* L. probably facilitated the shift from bee pollination to hummingbird pollination [36]. A study in *Silene littorea* discovered that a lack of anthocyanins exclusively in flower petals appeared as a stable polymorphism, but complete loss of anthocyanins in all plant parts was rare suggesting anthocyanins are the target of selection in photosynthetic tissues [37].

The flower pigmentation of *D. purpurea* is not uniform, but characterized by intensely pigmented dark spots on the bottom petals (**Fig. 1a**). Petal spots have been reported in other plant species including *Gorteria diffusa* [38], *Kohleria warszewiczii* [39], *Clarkia gracilis* [40], *Lilium leichtlinii* [41], and *Mimulus guttatus* [42]. Pigmentation spots on petals can be caused by the accumulation of high levels of anthocyanins (cyanidin derivatives) [40, 43]. These spots can be surrounded by a white region characterized by high levels of flavonols, the products of a pathway competing with anthocyanin biosynthesis [44]. Formation of intensely pigmented spots can be explained by a system involving a local R2R3-MYB activator (NEGAN) and a lateral R3-MYB repressor (RTO) [42]. NEGAN interacts with a bHLH and a WD40 partner (MBW complex) to activate *RTO* expression and RTO represses *NEGAN* expression by sequestering bHLH proteins required for the MBW complex [42]. RTO proteins appear to travel through the endosomal system to other cells, where they act as repressors [42]. Another study in *C. gracilis* reported an anthocyanin activating MYB (CgsMYB12) as the causal gene for the formation of a single large spot [45] indicating that alternative mechanisms have evolved to control the formation of pigmentation spots. Numerous functions have been described for such spots including an attraction and guidance of pollinating insects to increase the number of visits and to make each event more successful [39, 46–49].

The inflorescence of *D. purpurea* is usually constructed in the form of an indeterminate spike with zygomorphic tubular flowers on the side [50]. But in rather rare cases individuals develop large terminal flowers at the top of the spike, rendering the inflorescence determinate. These phenotypes show two types of flowers in the same plant: the normal zygomorphic flowers at the lower part of the spike and one actinomorphic structured flower at the top (**Fig. 2**). Keeble *et al*. investigated the heredity of this special trait and found it to be inherited recessively [51]. The characterization of the genetics behind the development of a terminal flower in *D. purpurea* has not been investigated yet. The occurrence of terminal flowers in plants with otherwise indeterminate inflorescence architecture has been described in different species like *Antirrhinum* [52] and Arabidopsis [53]. In *Antirrhinum*, a loss of function mutation in the *CEN* (*CENTRORADIALIS*) gene leads to the development of a terminal flower [52]. The functional *CEN* gene is expressed in the subapical meristem and disrupts the initiation of flower development [52]. Therefore, CEN is a flowering repressor. The *Arabidopsis* ortholog *TERMINAL FLOWERING 1* (*TFL1*) plays a similar role [54]. The mutant *tfl1* no longer represses flowering in the apical meristem, so it changes its fate and develops into flower primordia and subsequently into a flower [53, 55]. Both genes encode a protein belonging to the family of phosphatidylethanolamine binding proteins. Therefore, they are not conventional regulators of the transcription or translation of any specific target genes [56]. They have been shown to interact directly with other proteins so they might be part of a more elaborate signaling system [56].

**Fig. 2:**
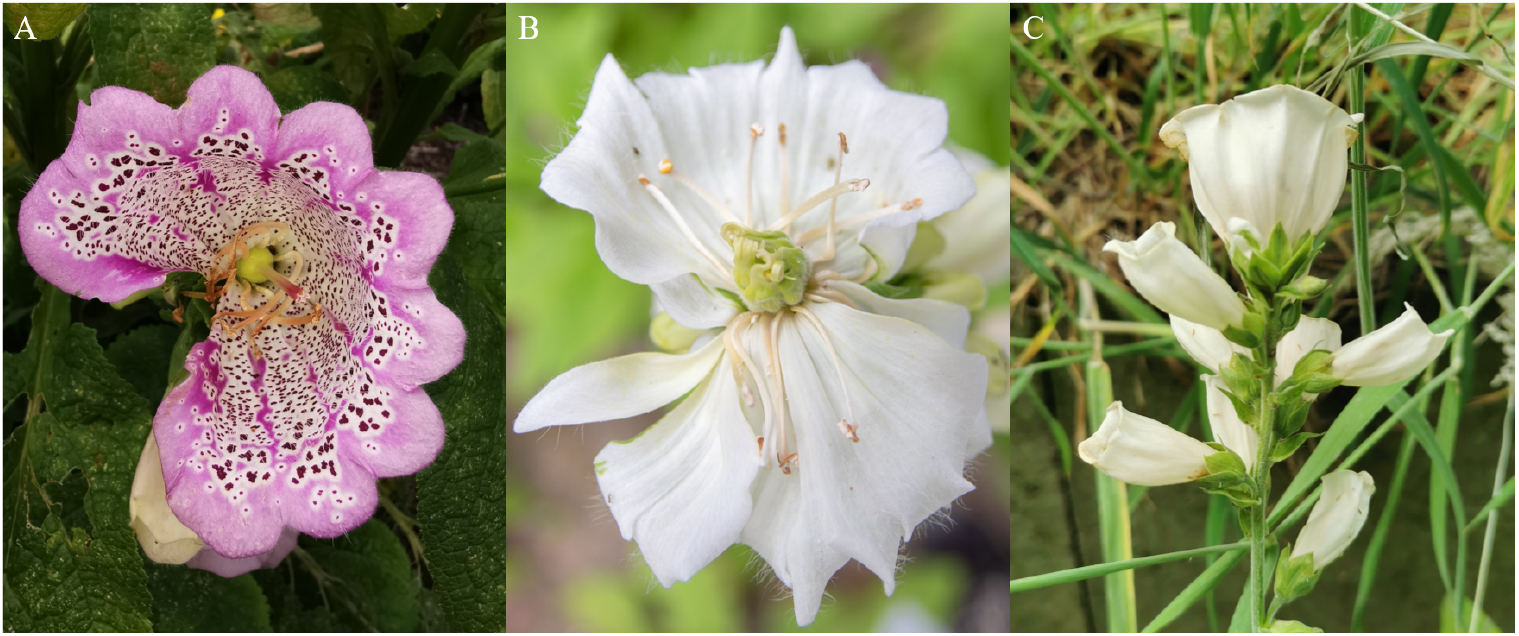
Three terminal flowers. (A): A magenta terminal flower with purple spots originated from the Botanical Garden of TU Braunschweig. (B and C): White terminal flowering plants.

Here, we report about the genome sequence of *D. purpurea* and the corresponding annotation. As a proof of concept, we demonstrate the power of this genome sequence for the discovery of specialized metabolism pathways by investigating the flavonoid biosynthesis. A comparison of white and magenta flowering individuals revealed molecular differences in the flavonoid biosynthesis genes that could explain the pigmentation loss in the white flowering plants. A candidate variant providing a possible explanation for large terminal flowers and indeterminate flowering architecture was discovered through comparative genomics based on this genome sequence.

## Materials & Methods

### Plant material and DNA extraction

*Digitalis purpurea* is a native species in Germany that is widespread at the borders of forests and in gardens. We collected seeds of a white and magenta flowering plant which were already in cultivation in a garden close to these coordinates 51.92351 N, 8.84441 E. No specific permissions were required. Formal identification of the plants was performed by Jakob Horz. Seeds of the investigated plants are now available in the Botanic Gardens of the University of Bonn (BONN-49855).

Plants were cultivated under long day conditions (16h light, 8h darkness) at 20°C. LEDs (Niello^®^, 300W) were used as the sole light source. Plants were incubated in the dark for two days immediately prior to the sampling of leaves for the DNA extraction to reduce polysaccharide content. Young leaves were harvested for the DNA extraction. The DNA extraction was performed with a CTAB-based protocol as previously described [57]. Quality and purity of the extracted DNA were assessed through NanoDrop measurement, agarose gel electrophoresis, and Qubit measurement. Short fragments (<25kb) were depleted with the Short Read Eliminator kit (Pacific Biosciences) (see [58] for a detailed workflow).

### Nanopore sequencing

The library preparation for sequencing was started with 1µg of high molecular weight DNA. The initial DNA repair was performed with two enzyme mixes of the NEBNext^®^ Companion Module Oxford Nanopore Technologies^®^ (ONT) Ligation Sequencing (E7180L) following the suppliers’ instructions. All following steps were performed according to the SQK-LSK109 protocol (ONT). Sequencing was performed with MinIONs using R9.4.1 flow cells. Upon blockage of a substantial number of nanopores, a wash step was performed with the EXP-WSH004 following ONT’s instructions to recover the blocked nanopores. Afterwards, a fresh library was loaded onto the flow cell. High accuracy basecalling was performed with guppy v6.4.6+ae70e8f (ONT) and minimap2 v2.24-r1122 [59] on a graphical processor unit (GPU) in the de.NBI cloud.

### De novo genome sequence assembly

A *de novo* genome sequence assembly was generated based on all long reads of a magenta flowering plant. NextDenovo v2.5.0 [60], Shasta 0.10.0 [61], and Flye 2.9.1-b1780 [62] were deployed to generate assemblies. Based on completeness, continuity, and assumed correctness, an assembly produced by NextDenovo v2.5.0 with read_cutoff = 1k, genome_size = 1g, and seed_depth = 30 was selected as the best representation of the genome. Additional polishing of this assembly was performed with NextPolish v1.4.1 [63] based on read mappings that were generated with minimap2 [59] and samtools v1.15.1 [64]. The genomic BUSCO assessment command contained the arguments ‘--long --limit 10 -sp arabidopsis’, 10^-3^ as e-value cutoff for BLAST, and eudicots_odb10 as the reference [65, 66]. AUGUSTUS v3.2.2 [67, 68] was used for the gene prediction as part of BUSCO. The assembly quality was further evaluated using k-mer analysis YAK (v0.1-r69-dirty) [69] (-k21) and Merqury v1.3 [70] with Meryl v1.4.1 (k=21) based on the ONT reads.

### Structural annotation

Paired-end RNA-seq reads (Additional file 1) were retrieved from the Sequence Read Archive via fastq-dump [71] and aligned to the assembled *D. purpurea* genome sequence with STAR v2.5.1b [72, 73] in 2-pass-mode using a minimal similarity of 95% and a minimal alignment length of 90% as previously described [74] to provide hints for the gene prediction process. Gene models were predicted with GeMoMa v1.9 [75, 76] using the following arguments: pc=true pgr=true p=true o=true GeMoMa.c=0.4 GeMoMa.Score=ReAlign Extractor.r=true GAF.f=“start==‘M’ and stop==^‘⋆’^ and (isNaN(score) or score/aa>=‘0.75’)”. In addition to RNA-seq hints, the annotation of the following datasets was supplied as hints: *Salvia hispanica* (GCF_023119035.1), *Rehmannia glutinosa* (GCA_016081115.2), *Sesamum indicum* (GCF_000512975.1), *Salvia miltiorrhiza* (GCF_028751815.1), *Penstemon davidsonii* (GCA_034814905.1), and *Penstemon smallii* (GCA_046254845.1). Predicted gene models were filtered via GeMoMa (f=“start==‘M’ and stop==^‘⋆’^ and (isNaN(tie) or tie>0) and tpc>0 and aa>50” atf=“tie>0 and tpc>0”) and renamed. BUSCO v3.0.2 [65, 77] was used to assess the completeness of the genome sequence and the completeness of the gene prediction. The completeness of predicted gene models was assessed with BUSCO v5.7.1 by running in proteome mode with default parameters.

### Functional annotation

Homologs in *Arabidopsis thaliana* were identified for the predicted peptide sequences to transfer annotation terms from this model organism to the *D. purpurea* sequences. Previously developed Python scripts [74, 78] were modified to enable the efficient identification of Reciprocal Best BLAST hits (RBHs) supplemented with best BLAST hits for query sequences that are not assigned to a RBH partner. BLASTp v2.13.0+ [79, 80] was run with an e-value cutoff of 0.0001. This approach allows an efficient identification of putative orthologs and forms the basis for the transfer of functional annotation details across species borders. *Arabidopsis thaliana* TAIR10/Araport11 annotation terms were automatically transferred to the predicted *D. purpurea* sequences. This workflow is implemented in the Python script construct_anno.py [81]. Enzyme and transporter encoding genes of the flavonoid biosynthesis were identified using KIPEs v3.2.6 with the flavonoid biosynthesis data set v3.4 [6, 82] with additional parameters listed in Additional file 2. Flavonoid biosynthesis controlling MYB transcription factors were annotated with the MYB_annotator v1.0.3 [83]. The following parameters were used for the MYB annotation: initial candidate identification via BLASTp v2.13.0+ [79, 80], alignment construction with MAFFT v7.475 [84], phylogenetic tree construction with FastTree v2.1.10 [85], minimal BLASTp alignment length of 75 amino acids, and 10 neighbors were used for the ingroup/outgroup classification (see Additional file 3 for details). The bHLH transcription factors were annotated using the bHLH_annotator v1.04 [86]. The following parameters were used for the bHLH annotation: identification of initial candidates via BLASTp v2.13.0+ [79, 80], alignment with muscle v5.1 [87], phylogenetic tree construction with FastTree v2 [85], a minimal BLASTp hit similarity of 40%, minimal BLASTp hit alignment length of 80 amino acids, and considering 10 neighbors for the ingroup/outgroup classification (see Additional file 4 for details).

### Annotation of transposable elements and repeats

Repetitive sequences were annotated with RepeatModeller2 [88] and RepeatMasker [89]. A species-specific repeat library was generated with RepeatModeller v2.0.7 using the assembled genome sequence as input. Genome-wide repeat masking and content assessment was then performed with RepeatMasker v4.2.2. with the parameters -e rmblast and -lib with the *Digitalis*-specific repeat library. EDTA v2.2.2 [90] was utilized for the genome wide identification and annotation of transposable elements. The parameters for the annotation process were chosen as follows: --species others --step all --overwrite 1 --sensitive 1 --anno 1 --threads 10 --force 1. For evaluating the annotation consistency the parameters were adjusted to: --species others --step anno --overwrite 0 --anno 1 --threads 10 --evaluation 1. For the annotation of TE diagnostic motifs, located on the inserted sequences in *ANS* and TFL1/*CEN*, first the sequence was translated based on putative open reading frames (ORFs) using getorf (EMBOSS v6.6.0.0) [91] with the parameter -minsize 150 to filter out short ORFs. Next the collection of putative polypeptide sequences was subjected to hmmscan (HMMER v3.4) [92] against the Pfam database [93]. The parameters used for hmmscan were --cut_ga and --noali. To find LTRs in the sequence inserted into the *ANS* locus, LTR_Finder v1.07 [94] was applied with default parameters. Two LTR sequences were extracted using bedtools getfasta v.2.31.1 [95] and subsequently aligned with MAFFT v7.525 [84]. The distance between the two sequences was calculated using distmat (EMBOSS v6.6.0.0) [91] with the parameter --nucmethod 2 in order to use Kimura’s 2 parameter model [96]. The evolutionary divergence and therefore insertion time of the TE can be estimated with the formula:

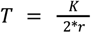

With T: Time, K: Substitutions per site and r: Neutral substitution rate.

As neutral substitution rates vary a lot and no values for closely related species of *Digitalis purpurea* are available, the calculation was performed with r = 1.5 X 10^-8^ a frequently used reported synonymous mutation rate in *Chalcone Synthase* and *Alcohol Dehydrogenase* loci of *A. thaliana* [97–99]. This approach is expected to return a rough estimation, while it is important to note that substitution rates can vary by over a magnitude [100].

As no LTRs were found in the sequence inserted into the *TFL1/CEN* locus, it was searched for possible poly(A) tracts, diagnostic for LINE and SINE non-LTR retrotransposition as reviewed by Kazuhiko Ohshima [101].

### Ks analysis and coverage analysis

To detect whole-genome duplication (WGD) events, a Ks analysis was performed. For each gene, the longest polypeptide sequence was selected. A self-BLAST of these polypeptide sequences was then performed using collect_best_BLAST_hits.py [81] with default parameters. The resulting BLAST table, the corresponding CDS file and a gene position file containing genomic coordinates were used as input for MCScan (via JCVI utility library 1.5.11, jcvi.compara.catalog) [102] to identify paralogous gene pairs in a synteny-aware manner. Codon-aware alignments of these gene pairs and calculation of Ks values were subsequently performed with ParaAT 2.0 [103] using the parameters -msa mafft, -format axt, -nogap and -kaks. The resulting ks distribution was plotted with a custom Python script and a Gaussian Mixture Model (GMM) with 5 components was fitted to the ks distribution using the GaussianMixture function from scitkit-learn [104].

To compare the coverage distribution of duplicated and single-copy BUSCO genes, the corresponding gene IDs were separated into two files and their genomic coordinates were extracted from the GFF annotation using the custom Python script extract_coordinates_from_gff.py [105]. Mean per-gene coverage was calculated using bedtools coverage v2.31.1 [95] with the -mean option. Coverage distributions were visualized as violin plots with box overlays using plot_coverage_busco2.py [105]. Peak detection was performed with SciPy’s gaussian_kde and find_peaks functions [106]. Statistical differences between the two distributions were assessed with a Mann-Whitney U test.

To assess the genomic heterozygosity, variants were called using bcftools v1.9 [107]. Raw alignment data were processed with bcftools mpileup using the parameters -q 20 (minimum mapping quality), -Q 15 (minimum base quality) and -a FORMAT/DP,FORMAT/AD to annotate read depth and allele depth. The resulting output was piped to bcftools call with the parameters --ploidy 2, -m (multiallelic caller) and -v (output variant sites only) to generate variant calls. High confidence heterozygous single nucleotide polymorphisms (SNPs) were extracted from the vcf file using bcftools view with the filters -i ‘GT=“het”’ and -v snps. The total number of heterozygous SNPs was counted to obtain the numerator for heterozygosity estimation. The number of callable genomic positions was determined using samtools depth with the same quality thresholds applied during variant calling (-q 20 and -Q15). Positions meeting the defined coverage depth criteria were counted to represent callable genome size. Relative heterozygosity was calculated as the ratio of heterozygous SNPs to the total number of callable genomic positions.

### Analysis of gene expression

Fastq-dump [71] was applied to retrieve the FASTQ files of *D. purpurea* RNA-seq experiments from the Sequence Read Archive (SRA). Gene expression was quantified using kallisto v0.44.0 [108] to process all RNA-seq data sets (Additional file 1) based on the predicted coding sequences. Individual count tables were merged with a customized Python script and served as the basis for a visualization of gene expression as previously described [17, 81].

### Identification of sequence variants associated with anthocyanin loss

Reads of two magenta flowering plants and two white flowering plants (Additional file 5) were aligned against the reference genome sequence of a magenta flowering plant with minimap v2.24-r1122 [59]. The resulting BAM files were indexed with samtools v1.13 [64]. Candidate genes associated with the flavonoid biosynthesis were manually inspected for systematic sequence differences via Integrative Genomics Viewer (IGV) v2.15.4 [109]. The same method was applied to one terminally flowering individual and three indeterminate flowering individuals.

### Identification of paralogous regions and functional assessment of the candidate *ANS* locus

To identify paralogous regions of the genome sequence in comparison to the contig harbouring *ANS* an alignment of ctg00580 against the genome assembly of *Digitalis purpurea* was performed using LAST v.1650 [110] with default parameters. The resulting mapping file was converted from MAF to TSV format using the maf-convert tab and filtered for alignments ≥ 2000 bp with percent identity ≥ 85%. Self-synteny analysis of the *D. purpurea* genome sequence was performed using JCVI v1.5.11. Synteny anchors were identified using jcvi.compara.catalog ortholog with --cscore=0.99 and --no_strip_names, and a microsynteny analysis for the *ANS* locus was conducted using jcvi.compara.synteny mcscan [102].

### Genotyping of a *Digitalis purpurea* population

A population of 89 plants was subjected to genotyping via PCR for the discovered mutations putatively associated with the white flower color or the changed flower morphology, respectively. The plants were first grown under long day conditions (16h light, 8h darkness) at 20°C. LEDs (Niello^®^, 300W) were used as the sole light source during that time. After four months the plants were moved outside the plant cultivation chamber and placed into two garden beds.

DNA for subsequent genotyping via PCR was extracted based on the GABI-Kat CTAB DNA preparation protocol [111] performed with one leaf from a single *D. purpurea* plant per sample. After extraction the DNA samples were stored at -20 °C.

PCR was conducted with oligonucleotides flanking the *TFL1*/*CEN* locus and a 106 bp deletion linked to the mutant *ans* allele at a distance of 1653 bp (Additional file 6, Additional file 7). Sizes of the PCR products were analyzed via agarose gel electrophoresis and revealed the genotypes.

Initial phenotypes of the plants resulting from the *ANS* mutation were derived from the visual inspection of the leaves as *D. purpurea* with a functional *ANS* would show magenta colored petioles (Additional file 8). These phenotypes were later validated by adding information about the flower color.

## Results

### Genome sequence assembly and structural annotation

A magenta flowering plant (DR1) was selected for sequencing and construction of a reference genome sequence [112]. The rationale behind this selection was the assumption that a magenta flowering plant should have all genes required for the biosynthesis of anthocyanins which are most likely responsible for the magenta flower pigmentation. For comparison, another magenta flowering plant (DR2) and two white flowering plants (DW1, DW2) were subjected to nanopore long read sequencing (Additional file 5).

Different assemblers were evaluated with respect to their performance in this study (**Table 1**). The assembly produced by NextDenovo2 was selected as the representative *D. purpurea* genome sequence due to the large assembly size of 940 Mbp, the high continuity (N50=4.3Mbp), and the high completeness of 96% of complete BUSCOs. Both quality values for the NextDenovo2 assembly estimated with YAK and Merqury resulted in 27.7 corresponding to ∼99.8 % base level accuracy.

**Table 1:** Comparison of assembler performance for the analysis of *Digitalis purpurea* long read sequencing data of the magenta flowering plant DR1.

| Assembly ID | DR1_v1 | DR1_v2 | DR1_v3 |
| --- | --- | --- | --- |
| Assembler | NextDenovo (v2.5.0) | Shasta (v0.10.0) | Flye (2.9.1-b1780) |
| Size | 940 Mbp | 773 Mbp | 1379 Mbp |
| # contigs | 684 | 4883 | 3738 |
| N50 | 4.3 Mbp | 0.9 Mbp | 1.9 Mbp |

| N90 | 464 kbp | 71 kbp | 132 kbp |
| --- | --- | --- | --- |
| BUSCO | C:95.7%<br>[S:63.5%,D:32.2%],<br>F:0.6%, M:3.7%<br>n:2326 | C:94.4%<br>[S:68.7%,D:25.7%],<br>F:0.6%, M:5.0%,<br>n:2326 | C:25.9%<br>[S:23.3%,D:2.6%],<br>F:6.0%, M:68.1%<br>n:2326 |

Only the representative genome sequence DR1_v1 was subjected to a gene prediction process that resulted in 35,916 protein encoding genes [113]. A completeness check revealed 96.3% complete BUSCO genes, of which a high proportion (55.8 %) were classified as duplicated. Comparison of mean read depth between single-copy and duplicated BUSCO genes revealed clearly distinct coverage distributions (Additional file 9 (Fig. S1)). Single-copy BUSCOs exhibited a single dominant peak at 31.8 x coverage, closely matching the average genome-wide coverage (33.6 x) (Additional file 9). In contrast, duplicated BUSCOs showed a bimodal coverage distribution, with one peak at 30.4 x and a second peak 14.8 x. The higher-coverage peak represents true duplicated loci (at merged haplophases), whereas the lower-coverage peak represents resolved haplophases. Together, these results indicate that the high proportion of duplicated BUSCO genes (55.8 %) can be explained by a combination of a recent WGD event and a partial haplotype separation in the assembly.

Ks analysis of syntenic paralog pairs identified a major peak at Ks = 0.21 and a less pronounced shoulder at Ks = 0.34, as inferred from Gaussian mixture modelling (**Fig. 3** and Additional File 9 (Fig. S2, Fig. S3)). Components at substantially higher Ks values were not considered further, as they likely reflect background noise rather than large scale duplication events [114]. Based on a Digitalis-specific repeat library, RepeatMasker estimated that interspersed repetitive sequences comprise 73.8 % of the genome sequence (Additional File 11). Annotation of transposable elements and repeats using EDTA revealed that TE derived sequences account for 67.4 % of the genome sequence. The majority of repeats are LTR retrotransposons which alone comprise 62.4 % of the assembly (Additional File 12). Genome-wide heterozygosity was estimated at 0.7 %, based on 2,028,683 heterozygous SNPs across 304,433,041 bp of callable sites in the genome sequence.

**Fig. 3:**
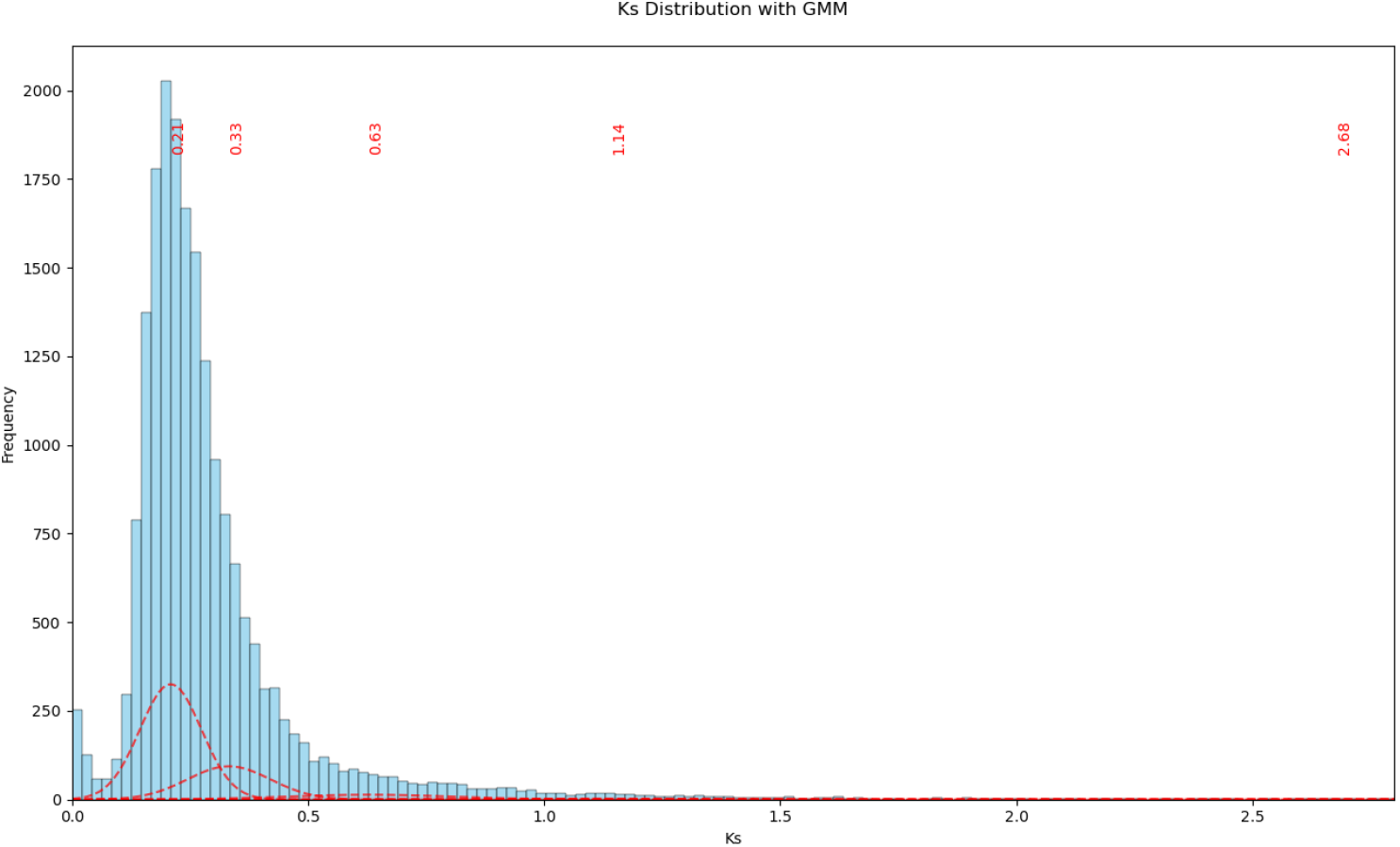
Ks distribution across syntenic gene pairs. Red dashed curves represent the individual components of a Gaussian mixture model (5 components) fitted to the Ks distribution, with red labels indicating the inferred Ks peak positions. The x-axis is limited to Ks ≤ 2.8 to improve visualization.

### Anthocyanin biosynthesis genes in *Digitalis purpurea*

Anthocyanins are well known pigments responsible for the coloration of many plant structures - especially flowers. Therefore, the genes encoding enzymes of the anthocyanin biosynthesis (*CHS, CHI, F3H, F3’H, DFR, ANS, arGST, 3GT*) were identified (**Fig. 4**; Additional file 13). Additionally, the transcription factors controlling the anthocyanin biosynthesis genes in *D. purpurea* were investigated. Genes belonging to the MYB75/PAP1 (DP112203, DP103545, DP109418) and MYB123/TT2 (DP102989) lineages were identified. In addition, a candidate for the bHLH partner in the anthocyanin biosynthesis MBW regulatory complex (bHLH42/TT8) was detected in *D. purpurea* (DP105472). An ortholog of the WD40 gene *TTG1* (DP111282) was also identified in the phylogenetic analysis. In summary, all genes expected for a complete anthocyanin biosynthesis pathway were successfully identified in the reference genome sequence DR1_v1.

**Fig. 4:**
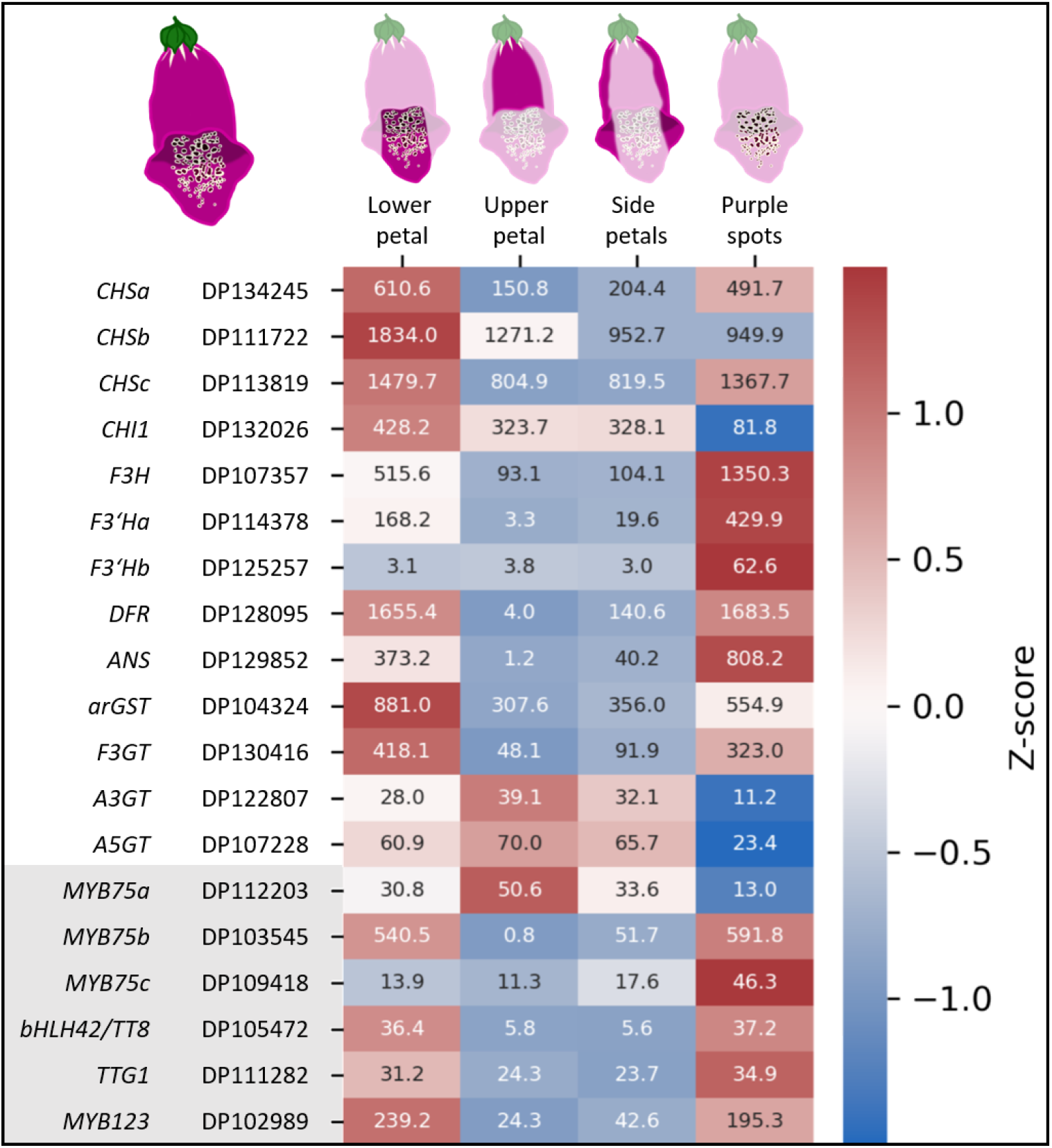
Expression plot showing the activity of anthocyanin biosynthesis associated genes across different petal tissues. The coloration indicates the z-score, the numbers inside the squares are the corresponding TPM-values. Abbreviations: *CHS*, chalcone synthase; *CHI*, chalcone isomerase; *F3H*, flavanone 3-hydroxylase; *F3’H*, flavonoid 3’-hydroxylase; *DFR*, dihydroflavonol 4-reductase; *ANS*, anthocyanidin synthase; *arGST*, anthocyanin-related glutathione S-transferase; *3GT*, anthocyanidin 3-O-glucosyltransferase; *F3GT*, UDP-glucose flavonoid 3-O-glucosyltransferase; *A3GT*, UDP-glucose anthocyanidin 3-O-glucosyltransferase; *A5GT*, UDP-glucose anthocyanidin 5-O-glucosyltransferase; MYB, Myeloblastosis; bHLH, basic helix-loop-helix; *TTG1, TRANSPARENT TESTA GLABRA 1*.

Additional support for candidate genes comes from a gene expression analysis that was conducted across a wide range of different samples (**Fig. 4**, Additional file 1, Additional File 14). Anthocyanin biosynthesis genes are expected to be highly expressed in pigmented structures like flowers and in particular within the intensely pigmented purple spots on the lower flower petal. A consideration of the gene expression pattern enabled a reduction of the candidate gene set to those gene copies which appear active in the anthocyanin-pigmented plant organs.

Genes encoding enzymes for all steps of the flavonoid and anthocyanin biosynthetic pathways were found to contain at least one paralog that is transcriptionally active in petal tissues. Two paralogous *F3’H* genes exhibited expression above baseline levels. *F3’Ha* (DP114378) was strongly active in floral tissues, whereas *F3’Hb* (DP125257) showed elevated expression primarily in root tissues (Additional File 14). Although roots typically do not accumulate anthocyanins, *F3’Hb* is reported here for completeness.

The putative anthocyanin-related transcription factor MYB75 is represented by three paralogous genes that displayed distinct expression profiles. *DpMYB75a* was active in the lower, upper, and side petal tissues but not in the spot sample. In contrast, *DpMYB75b* was strongly active in spot tissues and, consequently, also showed elevated expression in the lower petal samples, which include the spot region. *DpMYB75c* exhibited a comparatively uniform expression pattern across young floral tissues, with a modest increase in the spot samples.

### Sequence variants underlying pigmentation differences

All identified candidate genes associated with the anthocyanin biosynthesis were manually inspected in a read mapping comparing magenta flowering and white flowering plants. The most striking difference is an insertion of approximately 14.2 kb in the *ANS* gene (**Fig. 5**). This insertion is homozygous in the two white flowering plants, while the magenta flowering plants harbor one *ANS* allele with and one without this insertion and are therefore heterozygous (Additional file 15 (Table S3)). Although the reference genome sequence was derived from a magenta flowering plant, read mapping indicates that only the mutant allele is represented in the reference assembly, due to the heterozygous genotype of the individual. Since the reference genome sequence DR1_v1 contains the 14.2 kb sequence, the automatic prediction of the *ANS* gene model is inaccurate. The inserted sequence was misannotated as an intron of the *ANS* gene because no RNA-seq reads were aligned to the inserted region. However, in the wildtype allele, the first exon extends across the position of the insertion, indicating that the sequence cannot be removed by splicing in the mutant allele. This strongly suggests that the insertion disrupts the normal *ANS* transcript structure rather than representing an intronic region (**Fig. 5**). The inserted sequence was not annotated as a TE by EDTA. However, five Pfam domains located on five individual ORFs were found by hmmscan: zf-CCHC (Zinc knuckle), Pol_BBD (Pol polyprotein, beta-barrel domain), gag_pre_integrs (GAG-pre-integrase domain), SH3_retrovirus (Retroviral polymerase SH3-like domain) and RVT_2 (Reverse transcriptase) (Additional File 16). ORFs encoding RNase H domain and protease domains, essential for retrotransposons, were not found. Dating of the LTR sequences revealed a distance of 0.0265 substitutions per site. The inference of absolute insertion times with absent lineage-specific neutral substitution rates is imprecise. Nevertheless, LTR divergence has been widely used to infer relative TE age. With an assumed neutral substitution rate of 1.5 X 10^-8^ mutations per site per year, the insertion can be estimated to have taken place roughly 8.8 X10^5^ years ago. In the genomic region containing the transposable element (TE) insertion, the coverage plots of magenta-flowering plants do not display exactly half the read depth of adjacent genomic regions, as would be expected for heterozygous individuals (**Fig. 5**). Instead, the coverage is slightly elevated. Similarly, in white-flowering plants, the coverage across the insertion site is also somewhat higher than that of the surrounding sequences. These deviations from the expected pattern are likely caused by misaligned reads originating from additional copies of the TE elsewhere in the genome.

**Fig. 5:**
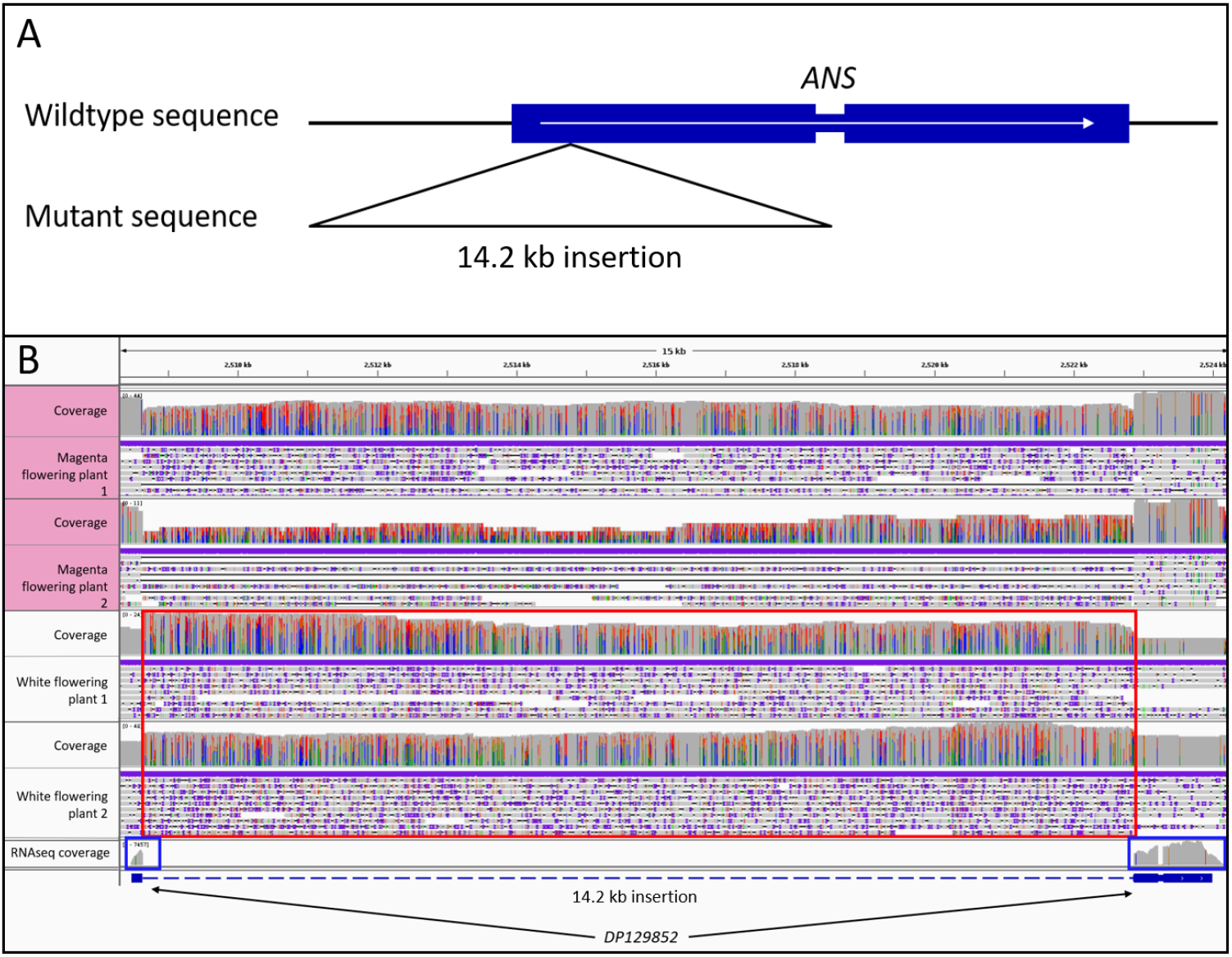
Comparison of the *ANS* locus in magenta and white flowering *D. purpurea* plants revealed a large insertion that appears homozygous in the white flowering plants. The insertion is expected to be a loss-of-function mutation that results in the loss of ANS enzymatic activity and thus a block in the anthocyanin biosynthesis ultimately resulting in the white flower phenotype. **A:** Schematic of the insertion. **B:** Screenshot from IGV with ONT read mappings of two magenta flowering plants and two white flowering ones at the *ANS* locus. The red box highlights the inserted reads. The blue boxes highlight RNA-seq read mapping coverage to corroborate the gene annotation.

To analyse the presence of additional *ANS* copies or homologous genomic regions resulting from the genome duplication, the contig harbouring *ANS* was aligned against the whole genome sequence. However, no alternative hits were identified (Additional file 17). While a microsynteny analysis revealed a syntenic region of the genes surrounding *ANS* on contig ctg000280, no *ANS* gene was identified on this contig (Additional file 18). This lack of detection in a highly continuous and complete genome sequence indicates that no alternative copy of *ANS* is present within the *D. purpurea* genome.

Genotyping of 89 plants with respect to the *ANS* locus identified 14 plants with the homozygous wildtype allele, 31 plants with a heterozygous genotype, and 44 plants with the homozygous mutant allele. In the genotyped population, 61 plants have flowered. The flowering phenotype was found to fit the genotype in 57 of these 61 plants, making up 93.4 %. Four plants showed a phenotype mismatching the genotyping results. Two plants flowered white while being heterozygous for the mutation and two plants showed magenta flowers but had a homozygous mutant genotype (Additional File 15 (Table S1)).

### Sequence variants leading to the formation of a large terminal flower

In the search for the genetic basis of the formation of large terminal flowers, the sequencing reads of one plant with a terminal flower were compared to three plants showing wildtype flower morphology. Upon investigation of the *Digitalis* orthologs of the *Arabidopsis TFL1* a 14.3 kb insertion was found starting only 4 base pairs downstream of the start codon of the gene and leading to a premature stop codon 4 codons further downstream (**Fig. 6**). The genotypes of the sequenced individuals at the *TFL1/CEN* locus can be found in Additional file 15 (Table S3). No LTR-sequences were found in the reassembled insert sequence. However, two significant hits for both the integrase domain (PF00665) and the Reverse Transcriptase domain (PF07727) were discovered by hmmscan. Also, a region of 131 bp containing two poly(A) sequences (50 and 48 bp in length, respectively) separated by 33 bp was found.

**Fig. 6.**
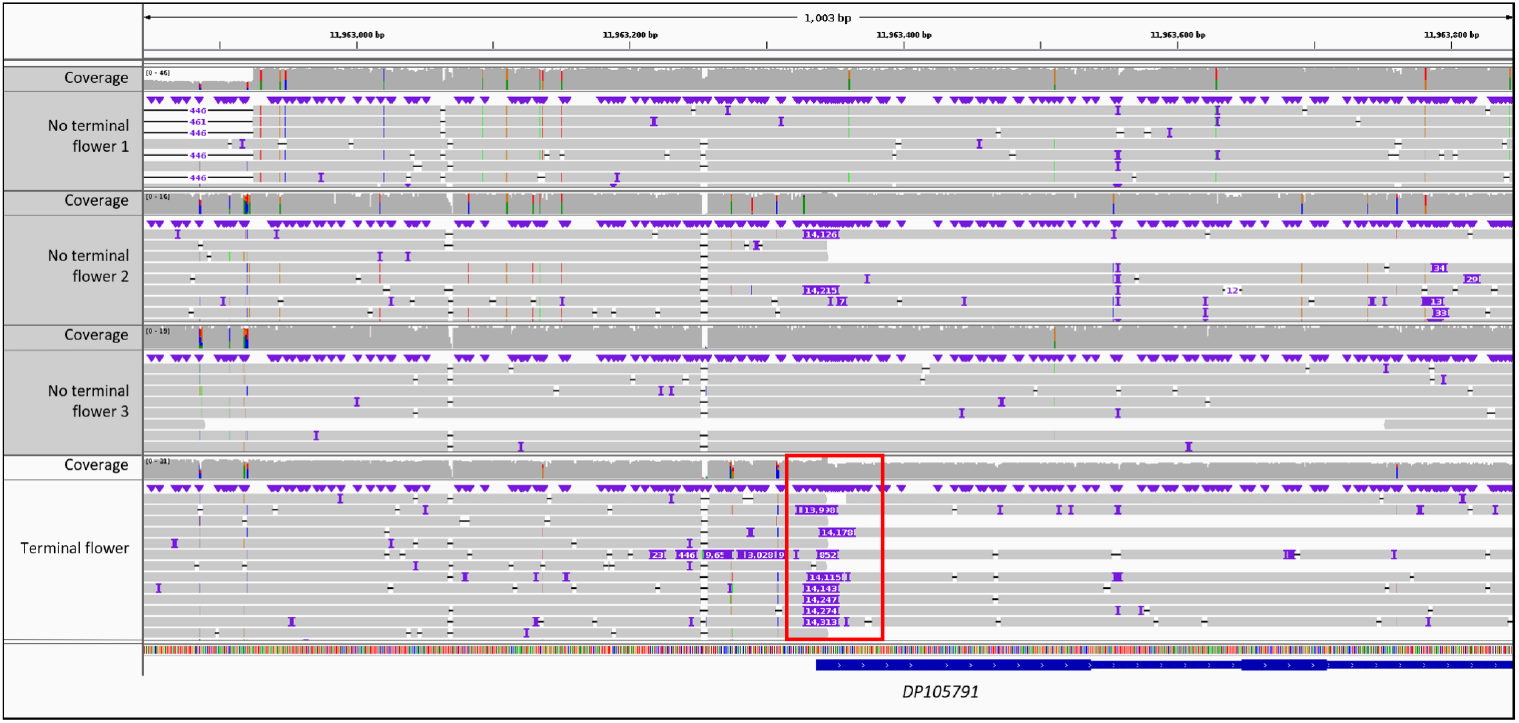
Genomic region showing the insertion of 14 kb in the *DpFL1/CEN* (DP105791) in the reads of the plant with a terminal flower (red box). The plants with indeterminate flowering architecture show at least some reads without the insert.

Out of 24 characterized plants, 23 showed the phenotype fitting the predicted genotype. While 17 plants showed the wildtype floral architecture and contained at least one wildtype allele, 7 plants developed terminal flowers. Of those plants with large terminal flowers, six had the homozygous mutant genotype and one appeared to have a heterozygous genotype (Additional File 15 (Table S2)).

## Discussion

The genome sequence of *D. purpurea* and the corresponding annotation of protein encoding genes enabled the identification of genes associated with the anthocyanin biosynthesis. Long read sequencing of two white flowering individuals enabled the discovery of sequence variants that might explain the lack of pigmentation. *ANS* encodes the anthocyanidin synthase, a crucial enzyme for the anthocyanin biosynthesis, and shows a homozygous insertion of a large DNA fragment in both white flowering plants, while the two magenta flowering plants show at least one allele without this insertion. The consequence of this insertion is most likely a loss of ANS activity that disrupts the anthocyanin biosynthesis. A loss of ANS activity fits all observations regarding the presence/absence of flavonoids in *D. purpurea*.

The insertion was found to contain TE-associated motifs like flanking LTRs as well as motifs of reverse transcriptase domain and GAG domain. Furthermore zinc knuckle, beta-barrel and SH3 retrovirus domains were detected. The lack of RNase H and protease domains combined with the fact that the detected motifs are not encoded by one continuous ORF but by five individual overlapping ORFs indicate that this TE is no longer functional and most likely heavily degenerated. This fits the obtained LTR dating results placing the insertion almost 900,000 years ago. When considering slower substitution rates available for plants, this time could increase drastically to well above 5 Mya [100]. Due to missing linear specific substitution rates the exact time remains unknown. It can be said that the insertion of the TE and thereby the occurrence of white flowering *Digitalis purpurea* plants predates the first horticultural efforts made by humans [115].

We noticed that flavonols are produced in the white flowering plants (Additional file 19). There is an overlap regarding the enzymes required for flavonol and anthocyanin biosynthesis. CHS, CHI, F3H, and F3’H are shared between both pathways. The presence of flavonols indicates that the genes *CHS, CHI, F3H*, and *F3’H* are functional, because they are required for the flavonol biosynthesis [116–120]. This observation reduces the set of remaining structural genes in the anthocyanin pathways to *DFR, ANS, arGST*, and *UFGT*. While we cannot completely rule out that *DFR, arGST*, or *UFGT* contribute to the loss of anthocyanin pigmentation, we have not observed any evidence for systematic sequence variants in these genes. Several previous studies discovered loss-of-function mutations in *ANS* as the cause of anthocyanin pigmentation loss [121–123]. The above described *ans* mutation could be a possible explanation for the lack of pigments in white *D. purpurea* flowers. Because the tested population did not show a perfect correlation between *ANS* genotype and floral phenotype, it remains possible that additional mechanisms contribute to the absence of pigmentation in white-flowering *D. purpurea* individuals. A further plausible explanation is that two plants grew in close proximity and physically merged, thereby compromising the representativeness of the sampled tissue. As transcription factors are often implicated in the loss of anthocyanin pigmentation [34], the identified candidate genes were checked for sequence variants associated with the anthocyanin loss. However, no variants fitting the phenotypes could be observed in these genes. Nevertheless, three paralogous *DpMYB75* candidates were found to have differential expression patterns in petal tissues. DpMYB75b is strongly active in the dark purple spots which suggests its involvement in the regulation of spot pigmentation DpMYB75a is expressed in the remaining floral samples but not in the spot sample. This suggests the existence of two separate regulatory complexes involved in the pigmentation of *Digitalis* flowers: One activating spot pigmentation (DpMYB75b) and one activating background pigmentation (DpMYB75a). Whether or not the absence of color and spot pigmentation in white flowering plants has any effect on pollination efficiency remains to be tested. However, there are many examples of plants where different visual cues in the flower lead to increased pollination efficiency [124–126]. The visual cues can consist of flower forms indicating landing sites for insects, simply pigmented flowers letting the flower stand out against the background or contrasting patterns indicating landing-position and -direction for insects on the flower [127, 128]. This suggests that the pigmentation patterns in *D. purpurea* flowers might contribute to increased attractiveness of the flower for pollinators. The contrasting spots on the lower petal possibly indicate a landing zone and the two UV fluorescent stripes at the base of the connation of the side petal with the lower petal may act as nectar guides (Additional file 19).

The present study focuses on explaining the completely white flowering phenotype (**Fig. 1**), but there are also cases of white flowers with magenta spots in *D. purpurea*. Similar patterns have been described before in other plant species [42]. It seems plausible to assume that a similar local activator/lateral repressor system comprising NEGAN and RTO also underlies the pigmentation pattern in *D. purpurea*. Further investigations are required to test this hypothesis.

*DpTFL1/CEN* is an ortholog of *AmCEN* and *AtTFL1*, genes crucial for the development of an indeterminate inflorescence architecture. Loss-of-function mutations of these genes have been shown to lead to large terminal flowers [52, 54]. The system responsible for inflorescence architecture seems to be highly conserved across plant species [56], suggesting a similar system in *D. purpurea*. Genotyping of a *D. purpurea* population for the *DpTFL1/CEN* locus revealed that plants exhibiting terminal flowers predominantly carry the homozygous mutant genotype (6/7), with one being heterozygous at the *DpTFL1/CEN* locus, suggesting possible mosaicism, misclassification or other influencing factors. If it is indeed only the homozygous mutant genotype that leads to the development of a determinate inflorescence architecture, this would corroborate the findings of Keeble *et al*. [51].

This study has two main limitations: (1) the number of sequenced individuals was limited (n=4), which constrains the ability to capture population-level genetic variation, and (2) PCR-based genotyping showed minor inconsistencies with the respective phenotypes (4 of 61 individuals for the *ANS* and 1 of 24 for the *TFL1/CEN* locus) indicating that the marker does not fully explain phenotypic variation or that other biological or technical factors contribute to mismatches. While the overall consistency was high, these factors should be considered when interpreting these results.

## Declarations

### Ethics approval and consent to participate

Not applicable

### Consent for publication

Not applicable

### Availability of data and materials

All analyzed RNA-seq data sets are publicly available from the Sequence Read Archive (SRA) (Additional file 1). ONT sequencing data of *Digitalis purpurea* are available from the European Nucleotide Archive (ENA) under PRJEB63449 (Additional file 5).The assembled genome sequence of DR1 and the corresponding annotation are available via bonndata: https://doi.org/10.60507/FK2/4KUKXI [113]. Custom Python scripts developed for this project are available via zenodo: https://doi.org/10.5281/zenodo.18637460.

### Authors’ contributions

BP planned the study. JH, KW, andRF performed the sequencing. JH and BP wrote the software and performed the bioinformatic analysis. JH performed genotyping experiments. JH, KW, RF, and BP interpreted the results, and wrote the manuscript. All authors have read the final version of the manuscript and approved its submission.

## Acknowledgements

This work was supported by the de.NBI Cloud within the German Network for Bioinformatics Infrastructure (de.NBI) and ELIXIR-DE (Forschungszentrum Jülich and W-de.NBI-001, W-de.NBI-004, W-de.NBI-008, W-de.NBI-010, W-de.NBI-013, W-de.NBI-014, W-de.NBI-016, W-de.NBI-022). We also thank all students participating in our ‘Data Literacy in Genome Research’ course and members of the research group Plant Biotechnology and Bioinformatics for discussion and support. We thank Hanna Marie Schilbert for spotting large terminal flowers on *Digitalis* and contributing to the brainstorming for experiments. We acknowledge support from Project DEAL and the University of Bonn for open access publication.

## Additional files

**Additional file 1**: *Digitalis purpurea* RNA-seq data sets that delivered hints for the gene prediction process.

**Additional file 2**: Documentation of all parameters and input files used for the annotation of the flavonoid biosynthesis genes with KIPEs3.

**Additional file 3**: Documentation of all parameters and input files used for the annotation of the MYB transcription factor gene family with the MYB_annotator.

**Additional file 4**: Documentation of all parameters and input files used for the annotation of the bHLH transcription factor gene family with the bHLH_annotator.

**Additional file 5**: *Digitalis purpurea* long read sequencing runs with corresponding accessions assigned by the European Nucleotide Archive (ENA).

**Additional file 6**: Design of the genotyping PCRs for the *ANS* and *TFL1/CEN* locus.

**Additional file 7**: PCR cycle program for the *DpCEN* and *DpANS* genotyping.

**Additional file 8**: *D. purpurea* plants showing magenta colored petioles.

**Additional file 9**: coverage plots of single copy and duplicated BUSCO genes; Ks plot of all syntenic gene pairs and Ks plot of all syntenic gene pairs in y log scale.

**Additional file 10**: Histogram showing the read depth across the entire assembly. A coverage of 20x suggests a successful separation of haplophases, while a coverage of 40x suggests that haplophases have been merged.

**Additional file 11**: Summary file output by RepeatMasker.

**Additional file 12**: Summary file output by EDTA.

**Additional file 13**: *Digitalis purpurea* candidate genes associated with the anthocyanin biosynthesis.

**Additional file 14**:Expression plot showing the activity of anthocyanin biosynthesis associated genes across different samples.

**Additional file 15**: Documentation of the genotyping results of a *D. purpurea* population at the loci *ANS* and *TFL1/CEN* and and the corresponding phenotypes as well as the genotypes of the four sequenced individuals.

**Additional file 16**: Map of the TE inserted into *ANS* depicting LTR sequences and hmmer motif-hits as well as their corresponding ORFs.

**Additional file 17**: Single LAST hit produced by searching the Assembly against the contig harboring the *ANS* gene.

**Additional file 18**: Microsynteny plot showing collinear relationships between the contig harboring the *ANS* gene and a detected syntenic region.

**Additional file 19**: Magenta flower (left) and white flower (right) under UV-illumination. The two upper petals were removed to open the flower and expose the inside of the remaining flower.

