## Additional file 3 for "Genome sequence of the ornamental plant *Digitalis purpurea* reveals the molecular basis of flower color and morphology variation"

MYB_annotator.py version: v1.0.3

MYB bait file: /vol/data/tools/MYB_annotator/MYB_baits.fasta	f92f35b0d351312c43a6e1b30aa0567e
MYB info file: /vol/data/tools/MYB_annotator/MYB_baits.txt	5836d7f542b672365883fa0a29d5d5a0
Subject FASTA file: /vol/data/20251010_Anno_files/DR1_v2.pep.fasta	6299a82bae835470942b2c9cc9a57837

Tool for initial candidate selection: blast
Tool for tree construction: fasttree
CPUs for BLASTp: 25
CPUs for RAxML: 25
Reference MYB file: AthRefMYBs.txt	abc8047bebc188b8a20935fe82467b5d
Type of input: PEP
blastp path: blastp
makeblastdb path: makeblastdb
mafft path: mafft
raxml path: raxml-ng
fasttree path: fasttree

Minimal BLASTp hit similarity cutoff: 60.0
Maximal number of BLASTp hits per bait: 100
Minimal BLASTp hit alignment length: 75
Number of neighbours to consider in classification: 10
Factor for branch length cutoff in identification of neighbours: 10.0
Minimal number of neighbours required for classification as MYB: 0
Distance cutoff for paralog clade masking: 10.0

MAFFT version: v7.505 (2022/Apr/10)
FastTree version:  2.1.11 
RAxML version: RAxML-NG v. 0.9.0 released on 20.05.2019 by The Exelixis Lab....
hmmsearch version detection failed.
