## Supplementary figures and images for "Genome sequence of the ornamental plant *Digitalis purpurea* reveals the molecular basis of flower color and morphology variation"

### Additional file 6

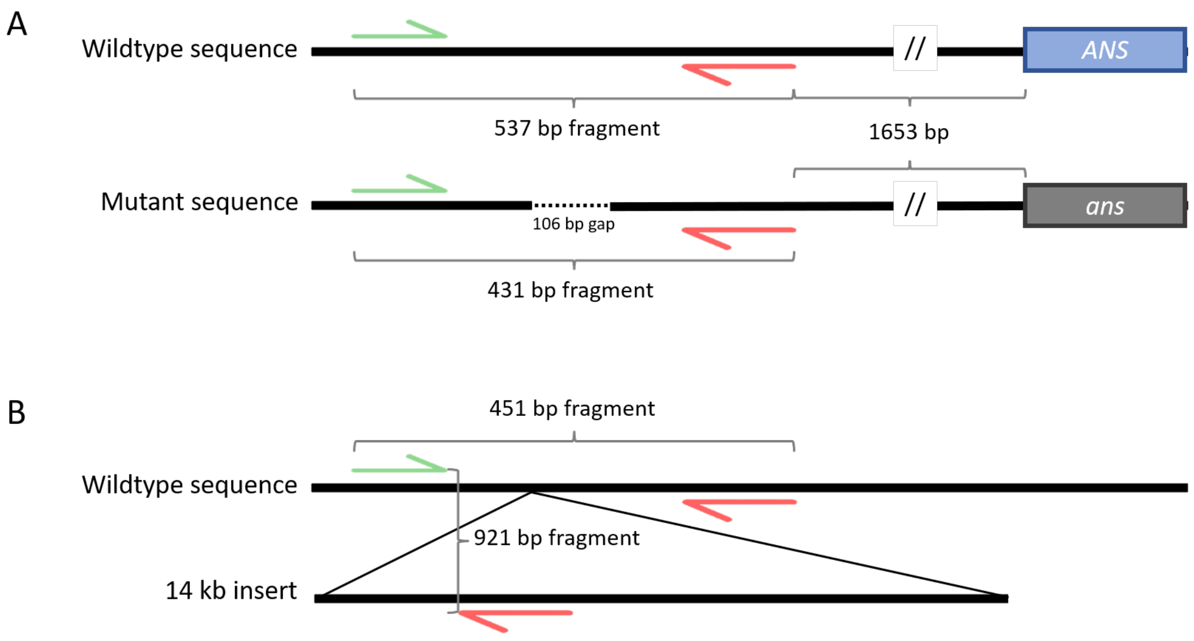


Design of the genotyping PCRs for the *DpANS* (A) and *DpTFL1/CEN* locus (B)

### Additional file 8

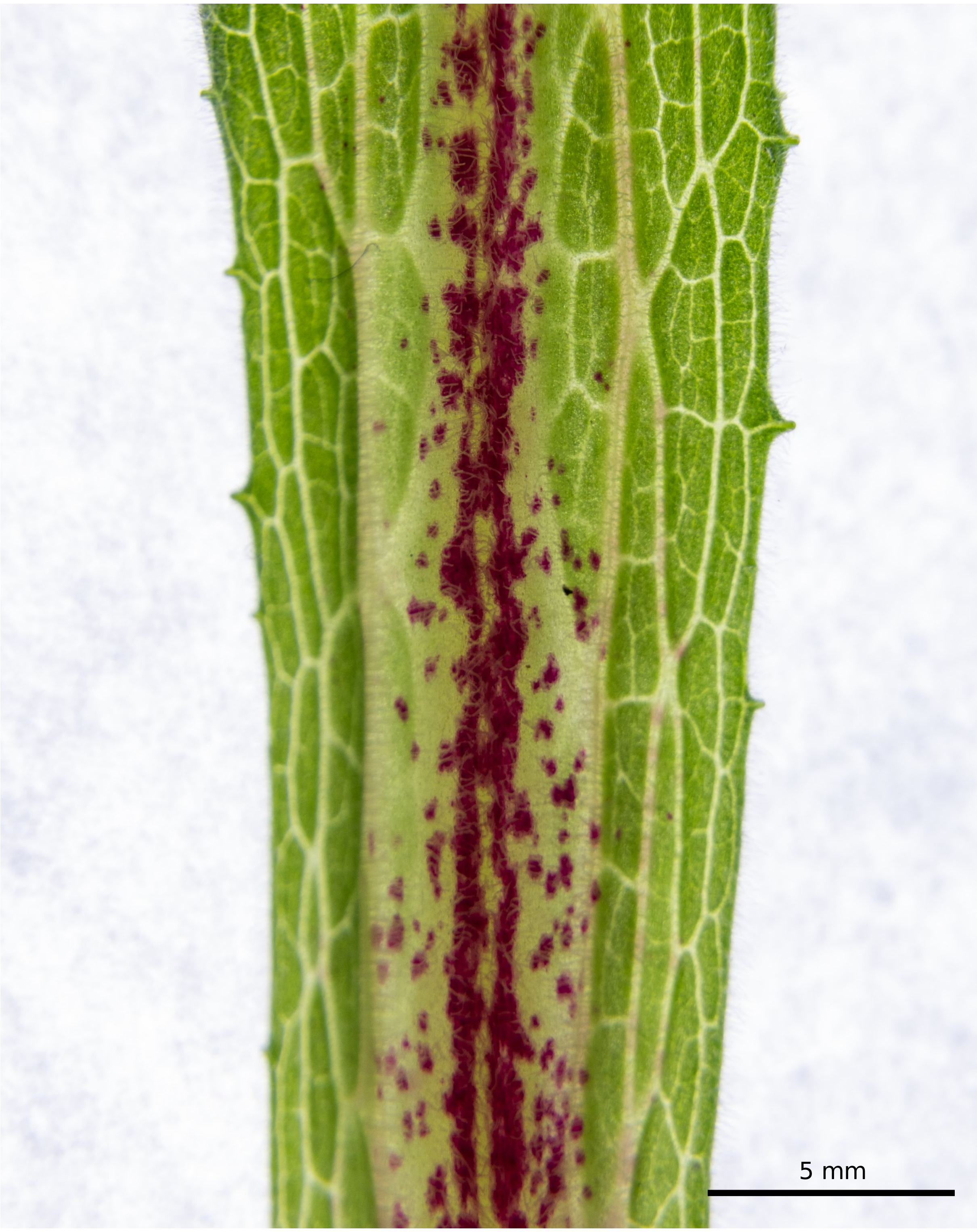


*Digitalis purpurea* leaf showing red pigmentation at the petiole.
