## Additional file 7 for "Genome sequence of the ornamental plant *Digitalis purpurea* reveals the molecular basis of flower color and morphology variation"

**TabS1**: PCR cycle program for the *DpTFL1/CEN* mutant. For the touchdown PCR program, the annealing temperature was decreased by 1 degree each cycle until the terminal temperature was reached. Number of cycles was 45.

| Step | Wildtype reaction | | Mutant reaction | |
|---|---|---|---|---|
| | Time | Temperature | Time | Temperature |
| Initial Denaturation | 5 min | 95 °C | 5 min | 95 °C |
| Denaturation | 1 min | 95 °C | 1 min | 95 °C |
| Annealing | 1 min | 65 °C -> 59 °C | 1 min | 65 °C -> 59 °C |
| Extension | 00:40 min | 68 °C | 1:05 min | 68 °C |
| Primers | JH1 | JH2 | JH2 | JH6 |

JH1 (wildtype reverse): 5’ AGGGTCTGTCATGATCTG 3’

JH2 (forward): 5’ TGTTCAATCCTCTTCATCTC 3’

JH6: (mutant reverse) 5’ TGTACGCGAATGAAGC 3’

**TabS2**: PCR cycle program for the *DpANS* mutant. Differently sized fragments will be obtained for wildtype and mutant target sequences, respectively. Number of cycles was 45.

| Step | Time | Temperature |
|---|---|---|
| Initial Denaturation | 5 min | 95 °C |
| Denaturation | 1 min | 95 °C |
| Annealing | 1 min | 59 °C |
| Extension | 00:45 min | 68 °C |

JH28: 5’ CTATAGTCGGGTCACATACGC 3’

JH29: 5’ CCTTTGAACATGGTGTCATAACC3’
