## Additional file 9 for "Genome sequence of the ornamental plant *Digitalis purpurea* reveals the molecular basis of flower color and morphology variation"

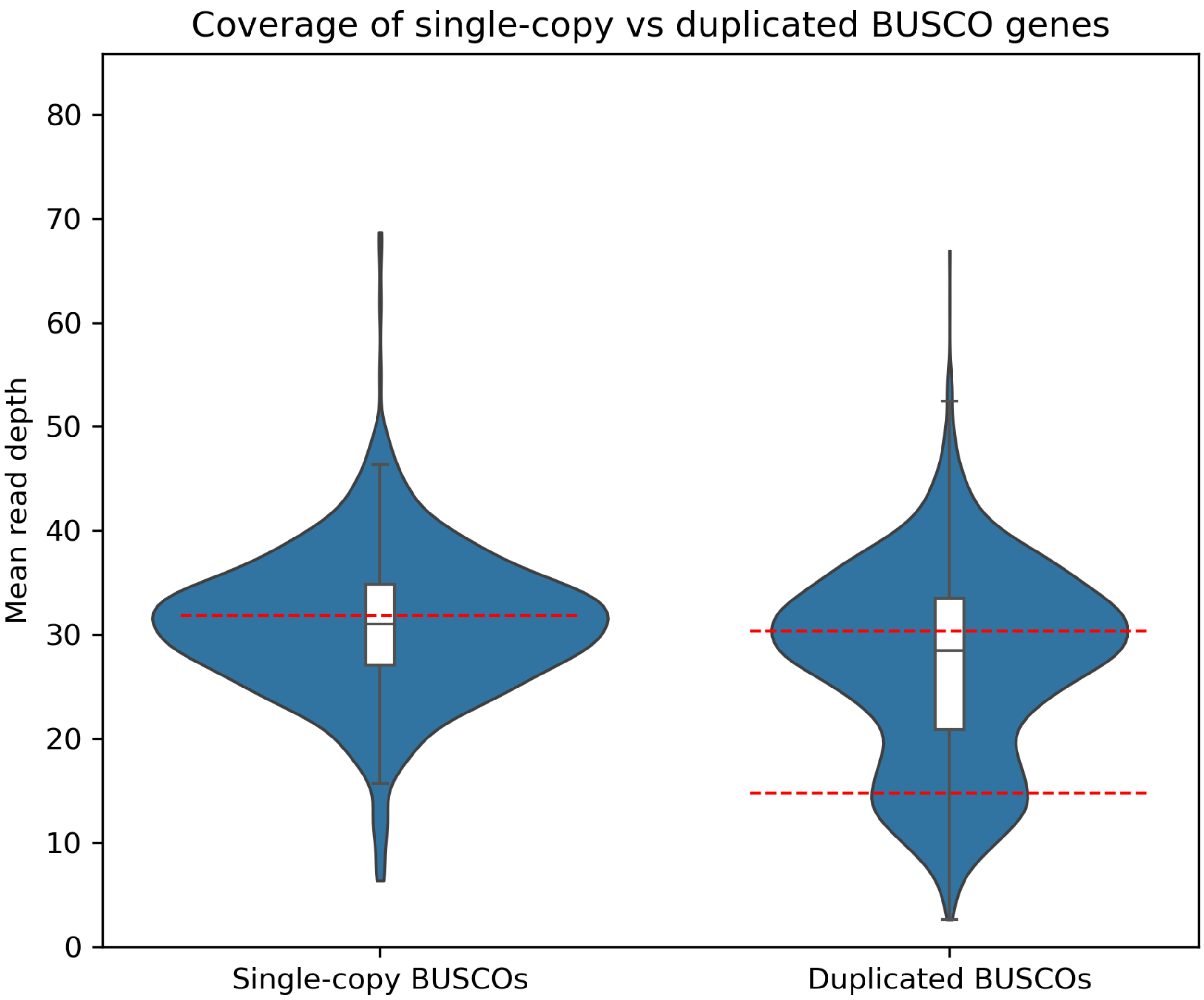


**Fig. S1:** Violin plots showing the distribution of mean read depth for single-copy and duplicated BUSCO genes. Red dashed horizontal lines denote coverage peaks identified by kernel density-based peak detection. Embedded box plots indicate median and interquartile range (IQR) with whiskers extending to 1.5x IQR.

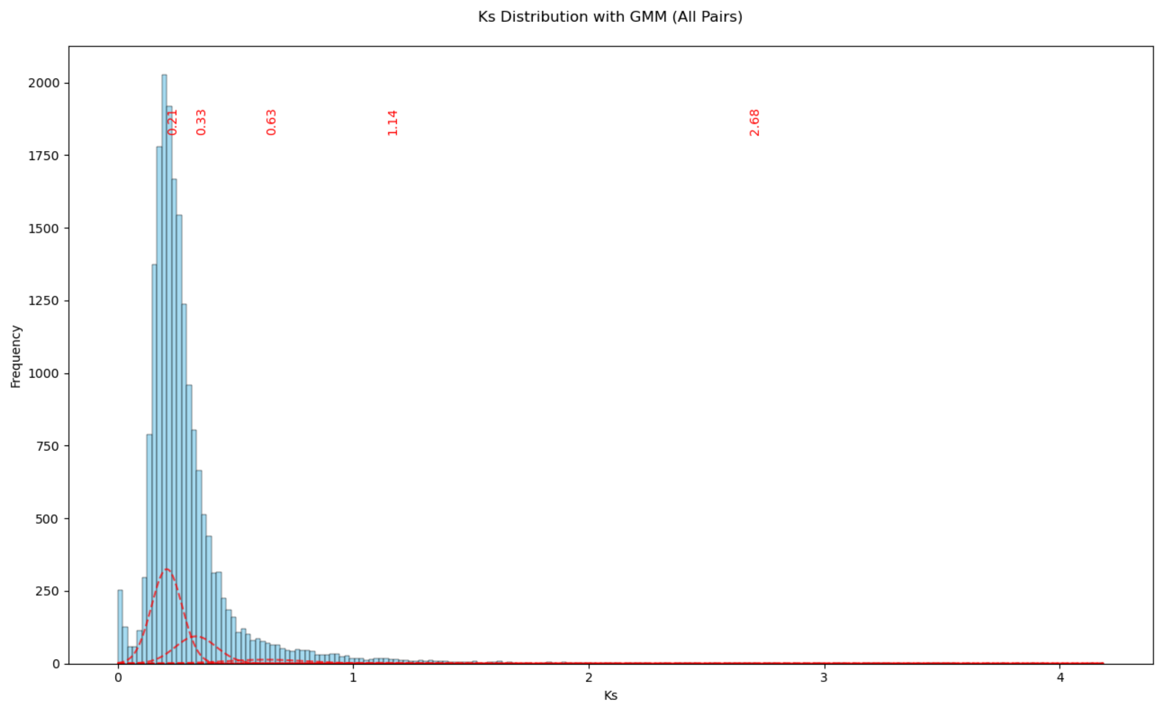


**Fig. S2:** Ks distribution across all syntenic gene pairs. Red dashed curves represent the individual components of a Gaussian mixture model (5 components) fitted to the Ks distribution, with red labels indicating the inferred Ks peak positions.

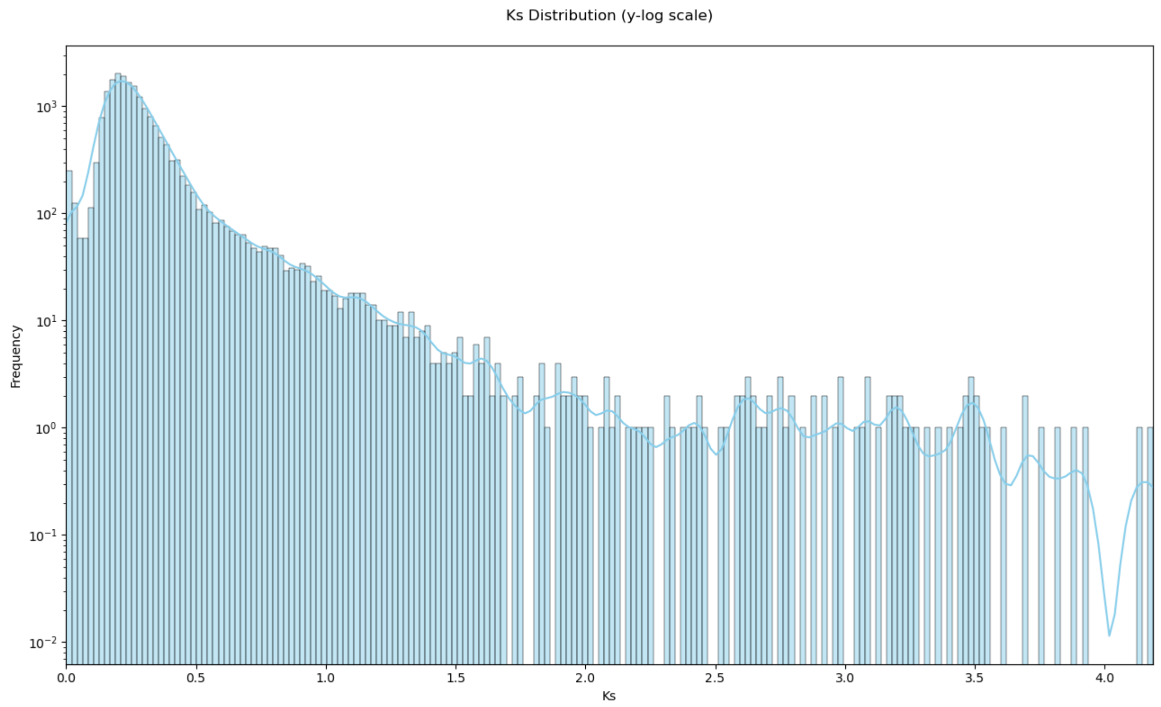


**Fig. S3:** Ks distribution across all syntenic gene pairs. Y-axis is displayed with log scale for visualization of lower occurrences
