## Additional file 10 for "Genome sequence of the ornamental plant *Digitalis purpurea* reveals the molecular basis of flower color and morphology variation"

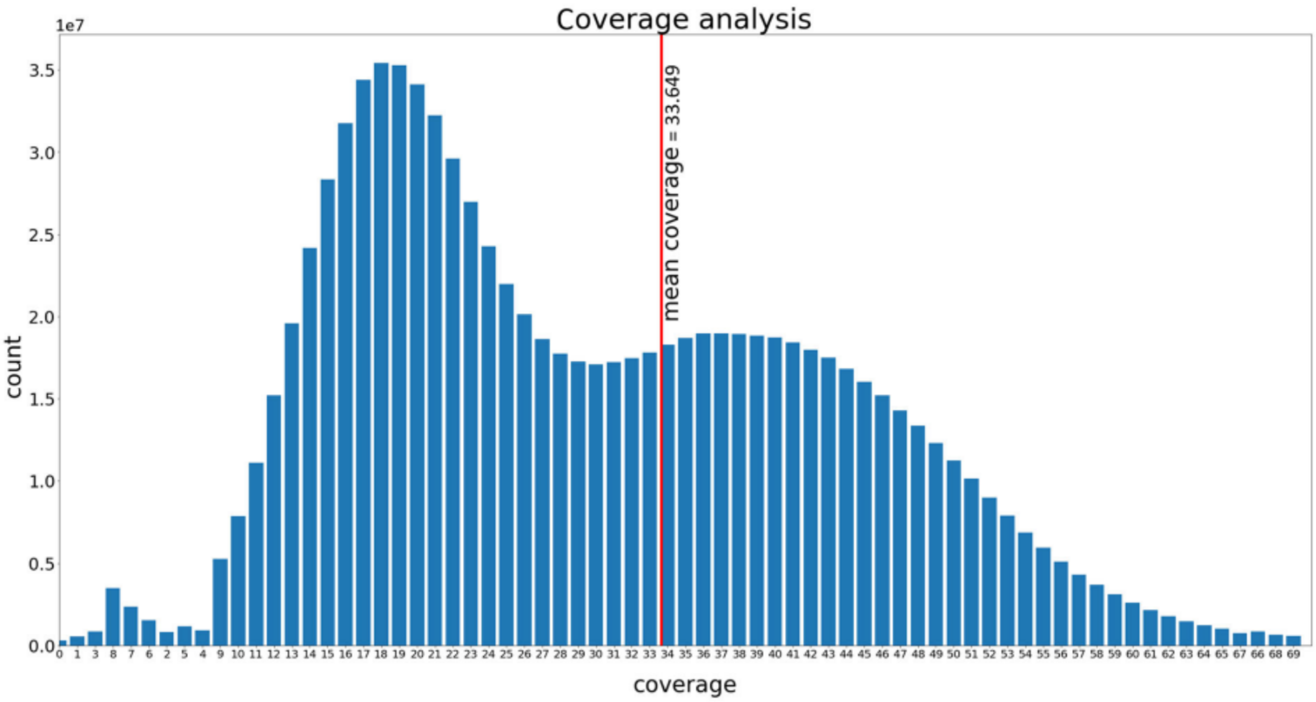


Coverage histogram of the DR1_v1 assembly. The coverage depth is displayed on the x-axis and the number of positions with a certain coverage is displayed on the y-axis. The mean coverage is depicted by the red line. The x-axis was cut at 69 for clarity.
