## Additional file 14 for "Genome sequence of the ornamental plant *Digitalis purpurea* reveals the molecular basis of flower color and morphology variation"

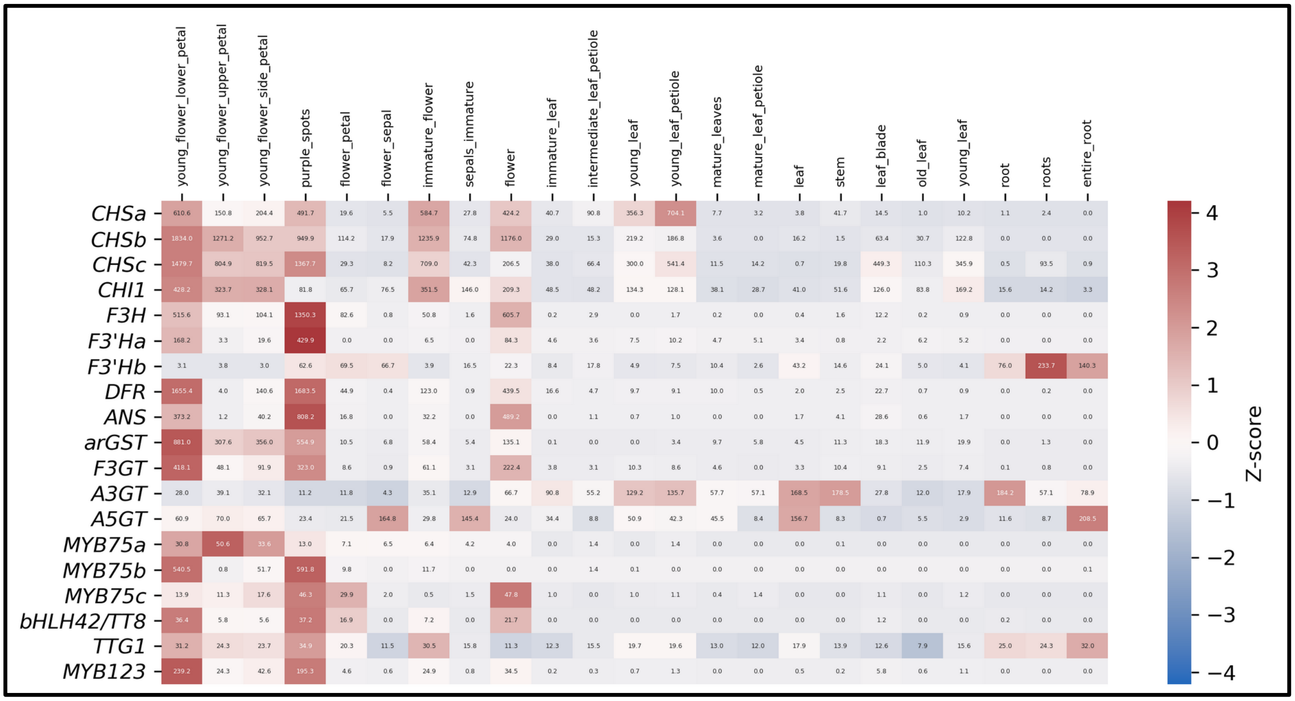


Expression plot showing the activity of anthocyanin biosynthesis associated genes across a range of different plant tissues. The coloration indicates the z-score, the numbers inside the squares are the corresponding TPM values. Abbreviations: *CHS*, chalcone synthase; *CHI*, chalcone isomerase; *F3H*, flavanone 3-hydroxylase; *F3’H*, flavonoid 3’-hydroxylase; *DFR*, dihydroflavonol 4-reductase; *ANS*, anthocyanidin synthase; *arGST*, anthocyanin-related glutathione S-transferase; *F3GT*, UDP-glucose flavonoid 3-O-glucosyltransferase; *A3GT*, UDP-glucose anthocyanidin 3-O-glucosyltransferase; *A5GT*, UDP-glucose anthocyanidin 5-O-glucosyltransferase; MYB, Myeloblastosis; bHLH, basic helix-loop-helix; *TTG1*, TRANSPARENT TESTA GLABRA 1.
