## Additional file 15 for "Genome sequence of the ornamental plant *Digitalis purpurea* reveals the molecular basis of flower color and morphology variation"

Table S1: Genotyping results at the *ANS* locus of 61 plants with their respective phenotype. A: Wildtype *ANS*, a: Mutant *ans*.

| Phenotype \ Genotype | A/A | A/a | a/a |
|---|---|---|---|
| Purple | 10 | 26 | 2 |
| White | 0 | 2 | 21 |

Table S2: Genotyping results at the *TFL1/CEN* locus of plants with their respective phenotype. T: Wildtype *TFL1/CEN*, t: Mutant *tfl1/cen.*

| Phenotype \ Genotype | T/T | T/t | t/t |
|---|---|---|---|
| Wildtype | 4 | 13 | 0 |
| Terminal flower | 0 | 1 | 6 |

Table S3: Genotypes of the sequenced individuals, derived from read mappings. A: Wildtype *ANS*, a: Mutant *ans*, T: Wildtype *TFL1/CEN*, t: Mutant *tfl1/cen.*

| ID \ Locus | ANS | TFL1/CEN |
|---|---|---|
| DR1 | A/a | T/T |
| DR2 | A/a | T/t |
| DW1 | a/a | T/T |
| DW2 | a/a | t/t |
