## Additional file 16 for "Genome sequence of the ornamental plant *Digitalis purpurea* reveals the molecular basis of flower color and morphology variation"

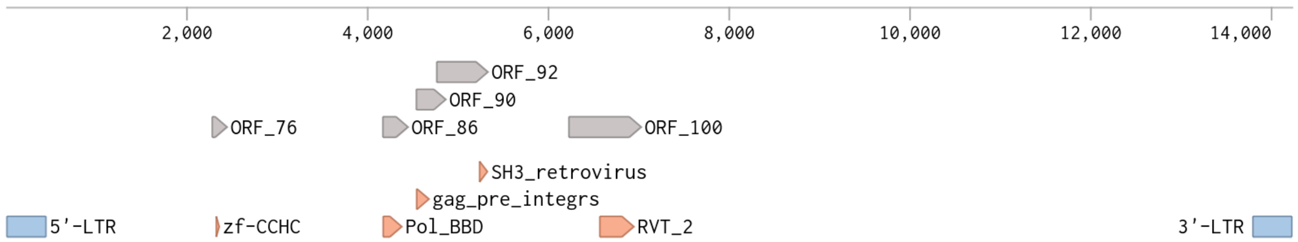


Elements on the putative TE inserted sequence in the *ans* gene. Blue: Flanking LTRs, Orange: Pfam hmmer motifs found on ORFs derived from the sequence. Gray: ORF for which the hmmer motivs were found. Sequence length is 14238 bp. ORF: Open reading frame, LTR: Long terminal repeat, zf-CCHC: zinc knuckle, Pol_BBD: Pol polyprotein, beta-barrel domain, gag_pre_integrs: GAG-pre-integrase, SH3_retrovirus: retroviral SH3-like fold, RVT_2: reverse transcriptase. Image was created with benchling.com
