## Additional file 18 for "Genome sequence of the ornamental plant *Digitalis purpurea* reveals the molecular basis of flower color and morphology variation"

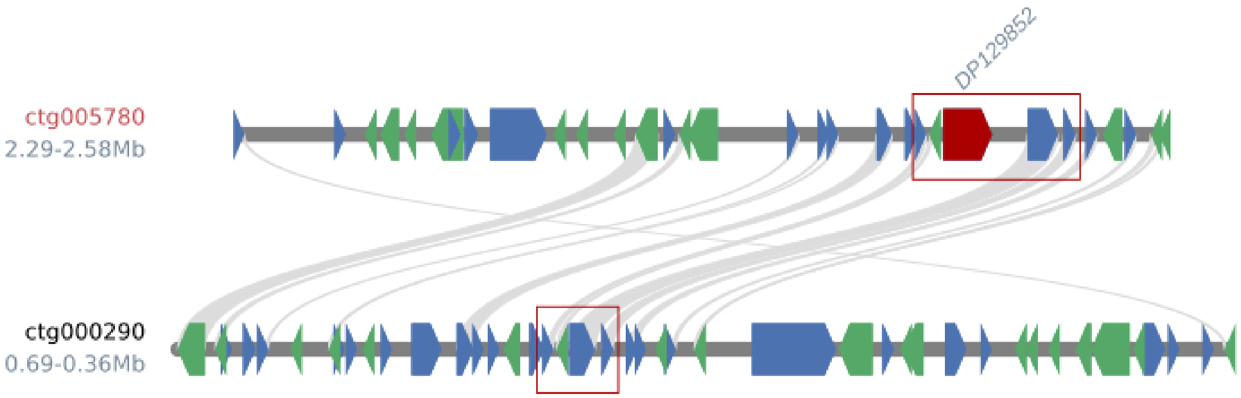


Microsynteny plot showing collinear relationships between the contig harbouring the *ANS* gene (highlighted in red) and a detected syntenic region (contig ctg000290). Red boxes highlight syntenic blocks comprising genes adjacent to ANS. No *ANS* paralog could be detected within the syntenic region on the paralogous contig.
