## Additional file 19 for "Genome sequence of the ornamental plant *Digitalis purpurea* reveals the molecular basis of flower color and morphology variation"

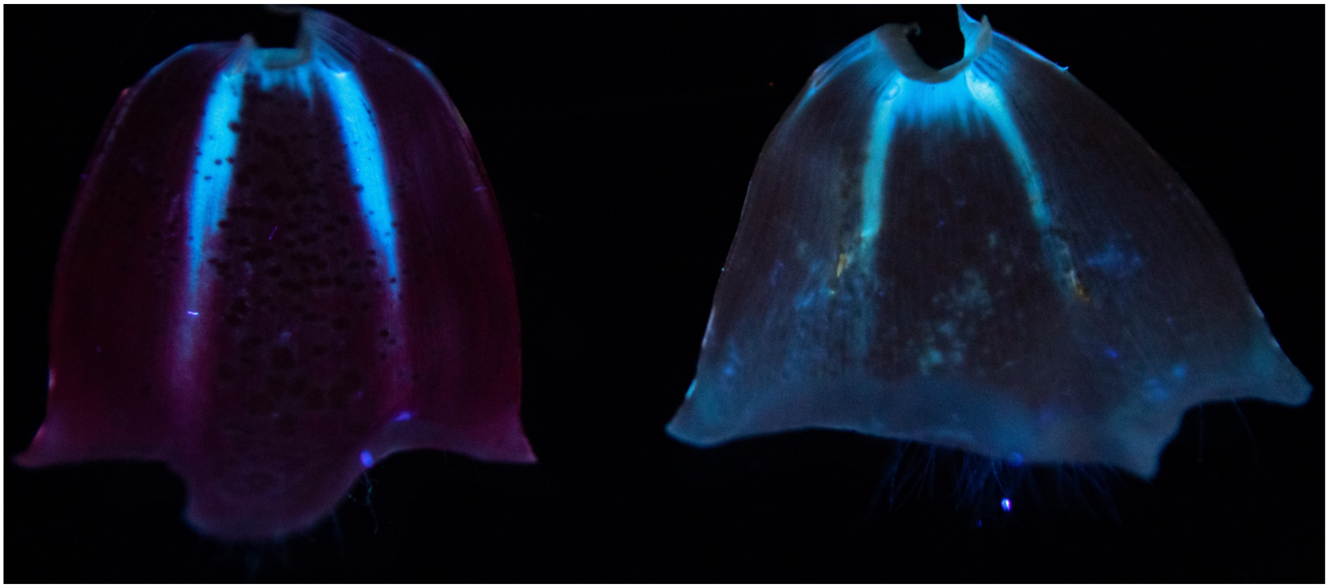

Magenta flower (left) and white flower (right) under UV-illumination. The two upper petals were removed to open the flower and expose the inside of the remaining flower.
